# BOLD-GPCRs: A Transformer-Powered App for Predicting Ligand Bioactivity and Mutational Effects Across Class A GPCRs

**DOI:** 10.1101/2025.08.04.668547

**Authors:** Davide Provasi, Kirill Konovalov, Nicholas Riina, Olivia Cullen, Marta Filizola

**Author notes:** Corresponding author: Marta Filizola, PhD, Department of Pharmacological Sciences, Icahn School of Medicine at Mount Sinai, One Gustave L. Levy Place, Box 1677, New York, NY 10029, USA.

## Abstract

G protein-coupled receptors (GPCRs) are important targets for drug discovery owing to their ability to respond to a broad range of stimuli and their involvement in numerous pathologies. Although traditional ligand-based and structure-based approaches have facilitated the development of effective therapeutics for many GPCRs, these approaches often fall short when applied to receptors with limited ligand or structural data. This limitation highlights the critical need for advanced strategies capable of accurately predicting ligand bioactivity across the entire GPCR family, especially for understudied receptor subtypes.

In this study, we introduce BOLD-GPCRs (BERT-Optimized Ligand Discovery for GPCRs), a deep learning framework designed to enhance the prediction of ligand bioactivity across class A GPCRs. Accessible via a user-friendly web interface, BOLD-GPCRs employs transfer learning and leverages curated datasets of known class A GPCR ligands, receptor sequences, and signaling-relevant mutations. By integrating dense neural network classifiers with transformer-based protein language models, BOLD-GPCRs captures complex relationships between receptor sequence/function and ligand activity. Our results demonstrate that BOLD-GPCRs achieves robust predictive performance for both ligand bioactivity and mutational effects across a broad range of class A GPCRs, underscoring its potential as a valuable tool for ligand discovery, especially for poorly characterized receptors.

**For Table of Contents Use Only:** *BOLD-GPCRs: A Transformer-Powered App for Predicting Ligand Bioactivity and Mutational Effects Across Class A GPCRs*

Davide Provasi, Kirill Konovalov, Nicholas Riina, Olivia Cullen, and Marta Filizola

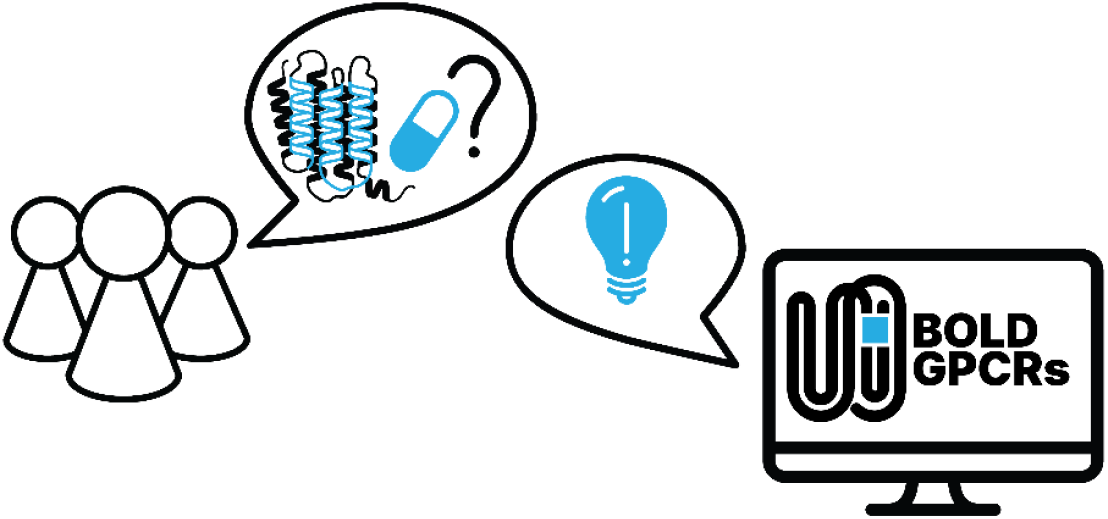

## INTRODUCTION

G Protein-Coupled Receptors (GPCRs)-mediated signaling pathways play a crucial role in regulating essential biological processes and diseases, including – but not limited to – development, cell survival, angiogenesis, and cancer. It is therefore unsurprising that approximately 35% of marketed drugs target GPCRs.^1^ However, only a small fraction of these receptors (134 out of ∼800, according to ref. ^1^) are targets for drugs approved in the United States or the European Union, leaving significant opportunities for the discovery of new GPCR-mediated therapies.

Deep learning architectures and other artificial intelligence (AI)-based methodologies hold strong potential to greatly accelerate the drug discovery process,^2, 3^ as demonstrated by their significant impact on recent campaigns.^4, 5^ By efficiently detecting patterns and deriving predictions from the analysis of large datasets,^6^ these models can improve the success rates of established computer-aided drug discovery strategies, such as ligand-based methods (i.e., those predicting biological activities based on features of known ligands) and structure-based techniques (i.e., those utilizing high-resolution protein structures for ligand design and optimization). Indeed, AI-driven approaches have already shown promise in GPCR drug discovery, for example in forecasting ligand-GPCR interactions, GPCR structures, novel GPCR ligands, and clinical responses.^7-9^ As the chemical, structural, and functional data on GPCRs continue to grow, these models will likely become even more powerful for uncovering new therapeutics.^10, 11^ A pressing question, however, is how to effectively leverage the currently available data to train models that can not only identify new bioactive ligands for GPCRs included in the training set but also generalize to less-studied receptors, such as orphan GPCRs, which lack robust chemical, structural, and functional information.

We have recently reported^12^ that advanced machine learning techniques, such as transfer learning,^13^ can address data scarcity at individual GPCRs by leveraging models pre-trained on larger datasets spanning entire GPCR subfamilies. This strategy is especially advantageous when training on both ligand-based and structure-based descriptors, as we have recently demonstrated for the opioid receptor subfamily.^12^ Specifically, transfer learning improved precision and recall for predicting bioactive opioid ligands across each opioid receptor subtype,^12^ whether we employed a dense neural network (DNN) architecture^14^ that integrates both structure-based and ligand-based information or a graph convolutional network (GCN) architecture^15^ relying solely on ligand-based descriptors.

To further enhance these deep learning architectures^12^ and accurately predict ligand bioactivities for GPCRs with limited data, we recently explored the synergy between these models and large language models (LLMs) designed for protein modeling^16, 17^ in predicting GPCR low-efficacy and biased agonists.^18^ Taking advantage of transformer architectures, LLMs such as BERT (Bidirectional Encoder Representations from Transformers)^19^ treat protein sequences as a “language”, identifying high-order relationships within sequences that inform protein structure and function. Indeed, the recently introduced ProteinBERT architecture was shown to effectively handle both classification and regression tasks with limited labeled data.^20^ This is achieved through self-supervised pre-training of a BERT model on the entire known protein space (∼106 million proteins derived from UniProtKB/UniRef90), and on Gene Ontology (GO) annotations of protein functions (∼45,000 terms spanning molecular functions, subcellular locations, and biological processes).^21^ A related approach, GPCR-BERT, used the ProtTrans architecture^22^ to predict GPCR sequence relationships and variations in conserved functional motifs with high accuracy.^23^

Here, we introduce BOLD-GPCRs (BERT-Optimized Ligand Discovery for GPCRs), a deep learning framework designed to accelerate ligand discovery for class A GPCRs, accessible through a user-friendly web interface. BOLD-GPCRs leverages transfer learning and integrates embeddings of class A GPCR sequences and functions via the ProteinBERT^20^ architecture with ligand embeddings based on physical, chemical, and topological properties. These protein and ligand representations are concatenated to form a simple multimodal embedding, which is then processed by a DNN to predict ligand bioactivity. Our results demonstrate that this enhanced approach accurately predicts ligand bioactivities across class A GPCRs, even in scenarios with limited chemical, structural, or functional data. Notably, by explicitly encoding residue-level information, the model can infer the effects of specific mutations on receptor activity and ligand binding. This capability is particularly valuable for elucidating the functional consequences of naturally occurring or engineered variants, guiding mutagenesis studies, and supporting drug design efforts targeting disease-associated mutant receptors. Overall, BOLD-GPCRs provides a valuable tool for discovering bioactive ligands for understudied or poorly characterized GPCRs, thereby advancing therapeutic development and precision pharmacology.

## METHODS

### Deep Learning Framework

We developed BOLD-GPCRs, a deep learning framework designed to enhance GPCR drug discovery using two embeddings: (i) a ligand embedding (depicted in purple in Figure 1), produced by a DNN that leverages molecular descriptors of the ligand and (ii) a GPCR embedding, obtained via the ProteinBERT transformer architecture,^20^ which processes both local features (i.e., sequences) and global annotations (i.e., functional and categorical annotations) of the receptor (shown in brown and green in Figure 1, respectively). These ligand and receptor representations are then fed into another DNN (highlighted in pink in Figure 1), which classifies each ligand-receptor pair as active or inactive based on whether the ligand has a reported bioactivity at that receptor with a potency below a chosen threshold (set here to 1 μM).

**Figure 1.**
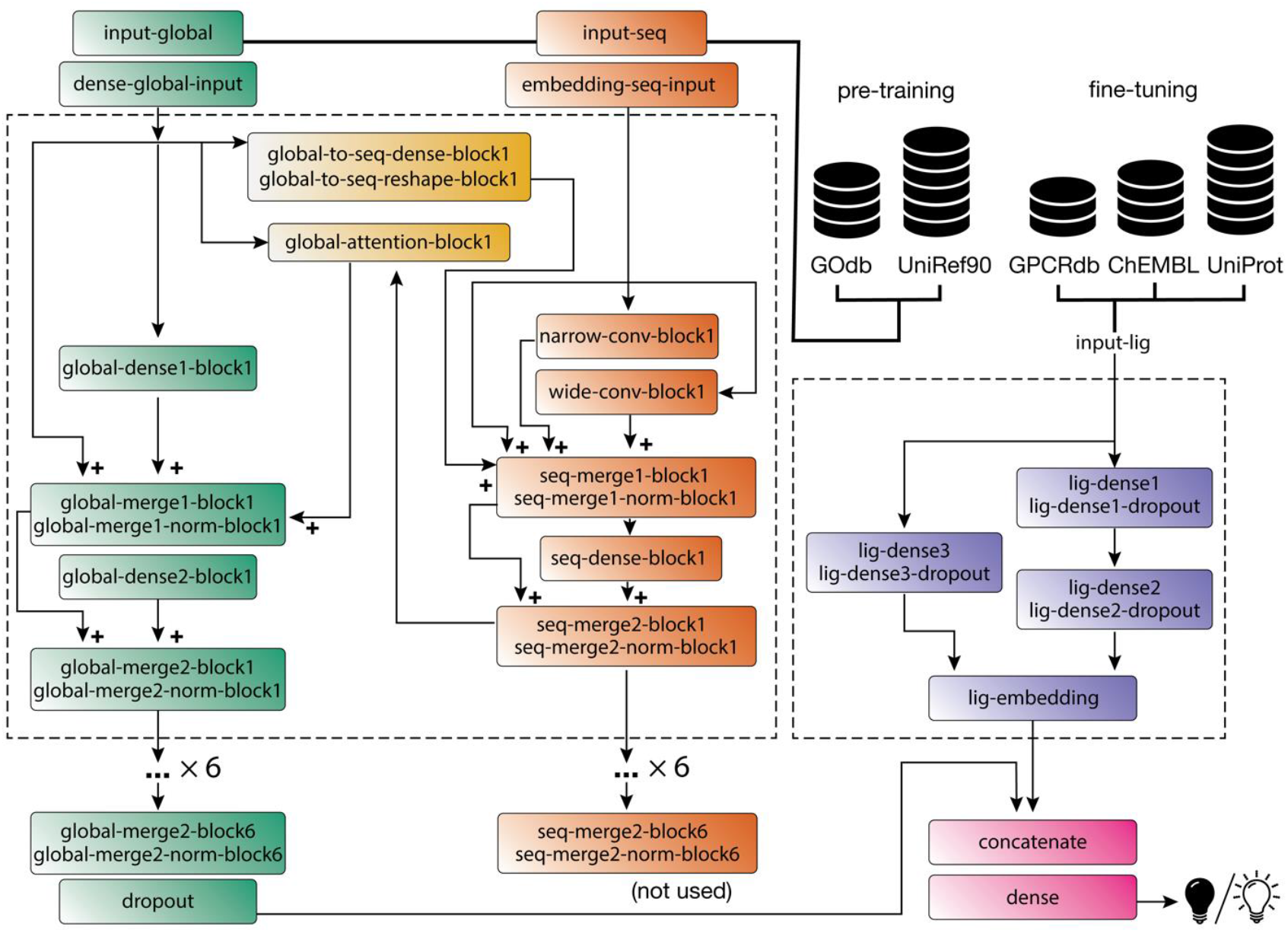
Schematic of the BOLD-GPCRs deep learning architecture. A DNN (in purple) generates an embedding of the ligand’s molecular properties retrieved from ChEMBL and GPCRdb, while ProteinBERT (in brown and green) produces an embedding of the receptor using a transformer architecture pre-trained on the entire known protein space, with sequences retrieved from UniProtKB/UniRef90 and functional annotations retrieved from GOdb. The two embeddings are then concatenated and fed into another DNN classifier.

For the ligand embeddings, ligand SMILES (Simplified Molecular-Input Line-Entry System)^24^ strings from the datasets described in the next section were converted to their canonical form using the CanonSmiles function in RDKit.^25^ The ComputeProperties function (also from RDKit) was then used to calculate 43 molecular descriptors, including basic physical and chemical properties, atom counts, and topological indices, as detailed in reference ^12^. These ligand descriptors were subsequently fed into two parallel functions (shown in purple in Figure 1): the first function comprised two dense layers with Rectified Linear Unit (ReLU) activation, each followed by a dropout layer, while the second function comprised a single dense layer with ReLU activation, also followed by a dropout layer.

For the protein embeddings, ProteinBERT version 1.0.1^20^ was employed. This model utilizes six transformer blocks operating through two interacting tracks to process both protein sequences (the local track, in green in Figure 1) and protein functions (the global track, in brown in Figure 1). Within each transformer block, fully connected layers link the global protein representation to the local representation. The local track processes the protein sequence through narrow and wide convolution blocks (with stride size of 1 and kernel size of 9 and 41, respectively), followed by dense layers (see Figure 1 for details). These local representations are then connected to the global representation through *global* attention layers (ochre in Figure 1). Specifically, each residue embedding *s*_*i*_ in the protein sequence is converted into a local key *k*_*i*_ = *σ*(*W*_*K*_*s*_*i*_), while the embedding of the global protein feature *x* is converted into a global query *q* = *σ*(*W*_*Q*_*x*), where *σ* is the Gaussian Error Linear Unit (GELU) activation function. Each key is scored as *z*_*i*_ = *ρ*(⟨*q, k*_*i*_⟩), where *ρ* denotes the soft-max function. An attention score is calculated by weighing the learned local values *v*_*i*_ = *σ*(*W*_*V*_ *s*_*i*_) by the score *z*_*i*_, yielding *y* = ∑_*i*_ *v*_*i*_*z*_*i*_. The attention score *z*_*i*_ indicates how strongly each residue contributes to updating the global protein embedding, thereby illuminating which residues are most relevant for predicting protein function. Finally, the ligand and protein embeddings are concatenated and passed through a final dense layer (in pink in Figure 1), which produces class probabilities via a soft-max activation function.

### Datasets and Approach for Training

We adopted a standard transfer learning workflow, starting with pre-training BOLD-GPCRs’ architecture on a larger protein dataset and subsequently fine-tuning it on a smaller set of non-olfactory class A GPCRs. Following the original ProteinBERT protocol,^20^ the BOLD-GPCRs’ architecture was first pre-trained on 106 million protein sequences from UniRef90, a non-redundant dataset of UniProtKB cluster representatives with at least 90% sequence identity. During pre-training, the 8,943 most frequent GO annotations associated with UniRef90 entries were used as global protein descriptors (capturing molecular functions, involvement in specific biological processes, and cellular localization) while the protein sequences provided local features. The loss function during self-supervised training was defined as the sum of the categorical cross-entropy for each amino acid position and the binary cross-entropy for these GO annotations.

For supervised fine-tuning, we used all non-olfactory class A GPCR data from ChEMBL^26^ and labeled ligands as active or inactive based on a potency threshold (“standard value”) of 1 μM. Specifically, every single-protein assay record with a “standard type” measurement (e.g., IC50, EC50, or AC50) was converted into a binary label (active/inactive) based on this 1 μM cutoff. This yielded 383,638 GPCR-ligand pairs, comprising 214,816 actives and 168,822 inactives (Table S1).

Wild-type GPCR sequences were retrieved from UniProt^27^ by cross-referencing target gene annotations with those in ChEMBL. To improve training efficiency, we excluded receptors with sequences longer than 600 residues, thereby removing targets such as LSHR and FSHR from the glycoprotein hormone receptors subfamily, TSHR from the thyrotropin-releasing hormone receptors subfamily, and RXFP1 from the relaxin family peptide receptors subfamily. Table S1 summarizes the total number of ligands, receptors, and bioactivity datapoints, while Table S2 shows their distribution across receptor subfamilies. Data on receptor mutation effects were sourced from GPCRdb.^28^ Only entries with specified ligands and quantified potency changes were retained. The final mutation dataset contained 10,920 datapoints, encompassing 592 unique ligands, 2,459 unique mutations, and 45 distinct targets. Notably, 93% of these datapoints involved active ligand–receptor pairs. Supervised fine-tuning was carried out in two stages: first, training for 10 epochs with the ProteinBERT layers frozen, and then training for 10 epochs with all model parameters unfrozen. Binary cross-entropy was used as the loss function throughout training.

We conducted 5-fold splits (using 80% of the data for training and 20% for test/validation) across three distinct scenarios: (i) a “random split”, where individual ligand-target records were randomly assigned to either the training or validation sets; (ii) a “ligand split”, where all records associated with a given ChEMBL molecule ID were grouped entirely into either the training or validation set; (iii) a “target split”, where all records associated with a given ChEMBL target ID were assigned exclusively to one of the two sets.

### Baseline Models

To assess the performance gains from the LLM architecture, we compared it with two simpler baseline models: a DNN and a random forest classifier. The DNN closely mirrored the ligand embedding network in architecture, activation functions, and optimization algorithm, but it was trained on a concatenated vector that combined the same ligand descriptors with a one-hot encoding of the target. The random forest classifier was trained on this same input vector.

### Effective Training Size and Heterogeneity of the Dataset

To quantify the amount of relevant training data available for each GPCR, we measured pairwise similarity between targets using two complementary metrics: one based on sequence similarity (assessed by PSI-BLAST scoring of pairwise structural alignments^29^), and another based on pharmacology similarity [assessed by the Similarity Ensemble Approach (SEA),^30^ which relates proteins based on the set-wise chemical similarity among their ligands].

To compare receptor sequences, we recovered pairwise structural alignments from GPCRdb^28^ that incorporate information from experimentally determined high-resolution GPCR structures. We computed a raw alignment score *S* using the BLOSUM-62 amino acid substitution matrix^31^ with affine gap costs^32^ of *k* + 10 for a gap of length *k*. We then normalized *S* as follows:

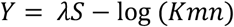

where *m* is the size of the comparison database, *n* is the length of the aligned sequences, and *λ* = 0.252 and *K* = 0.035 are standard parameters for the BLOSUM-62 matrix.^33^ Assuming that *y* follows a standard Gumbel distribution with location 0 and scale 1, the probability of observing a raw score higher than *S* by chance (the p-score) is:

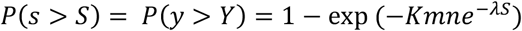

The quantity *E* = *Kmne*^−*λS*^, commonly reported in PSI-BLAST searches, measures the alignment’s significance. When E is small, *e*^−*E*^ ≈ 1 and *P*(*s* > *S*) ≈ 0, indicating that obtaining a score better than *S* by chance is unlikely, and thus the two sequences are likely related. In contrast, large positive *E* values imply *e*^−*E*^ ≈ 0 and *P*(*s* > *S*) ≈ 1, suggesting that the alignment may be due to chance.^34^

To compare GPCRs based on the similarity of their ligands, we used the metric introduced in SEA.^35^ Specifically, we computed a raw score *S* = ∑ *T*_*ij*_*H*(*T*_*C*_ − *T*_*ij*_) where *T*_*ij*_ is the pairwise Tanimoto coefficient between two ligands, *H* is the Heaviside step function, and *T*_*C*_ is a Tanimoto threshold set to 0.57 following reference ^30^. The score was then standardized to *Z* = (*S* − *μ*)/*σ* where *μ* and *σ* are functions fitted to the sizes of the respective ligand sets. Following reference ^30^, we treated the score Z as the standardized variable of a Gumbel distribution *y*∼Gumbel(0,1), i.e.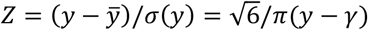, where *γ* is the Euler-Mascheroni constant (the mean of *y*). The p-score is computed as:

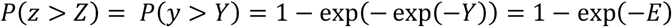

where 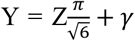 and 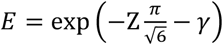, with small values of *E* corresponding to *P*(*z* > *Z*) ≈ 0 (see reference ^35^ for additional technical details).

For a given GPCR *j*, we defined its effective training size *n*_*i*_ by summing the training set sizes of all targets *i* that are sufficiently similar to *j*, i.e., those for which *E*_*ij*_ < *E*_*c*_ where *E*_*ij*_ is either the PSI-BLAST or SEA similarity between *i* and *j*, and *E*_*c*_ is a similarity cutoff value. Formally,

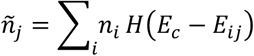

where *H* is again the Heaviside step function. We then calculated the active/inactive imbalance in the effective training data for GPCR *j* as:

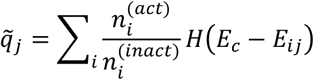

where 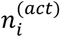 and 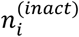 are the numbers of active and inactive ligands, respectively, in the training set of GPCR *i*. If 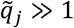, the effective training set for *j* is dominated by active ligands; if 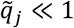, it is dominated by inactive ligands.

Lastly, we assessed the effective training set heterogeneity for GPCR *j* by calculating the average pairwise Tanimoto coefficient for all ligand pairs (*a, b*) of each GPCR *i*, and summing over all targets similar to *j*:

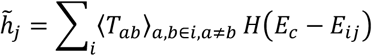

A value of 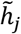 close to 1 indicates that the ligands in *j*’s effective training set are highly similar to one another (i.e., low heterogeneity), whereas lower values suggest greater chemical diversity in the effective training set. By examining the effective training set size *ñ*_*j*_, active/inactive imbalance 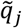, and heterogeneity 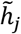 together, we obtained a comprehensive view of the relevant and diverse training data available for each class A GPCR.

## RESULTS

### BOLD-GPCRs’ Deep Learning Architecture Outperforms Classical DNN and Random Forest Models in Predicting the Bioactivity of All Class A GPCR Ligands

We first assessed the performance of the BOLD-GPCRs’ deep learning architecture illustrated in Figure 1 by training on the complete collection of ligands, receptors, and bioactivity data available for Class A GPCRs (Table S1). On a random test set representing 20% of the Class A GPCR data, the BOLD-GPCRs’ architecture achieves an average recall of 0.85 and an average precision of 0.72 (Figure 2). Confidence intervals across five different splits of the data into training and test sets are reported as the 25^th^ and 75^th^ quantiles in Table S3.

**Figure 2.**
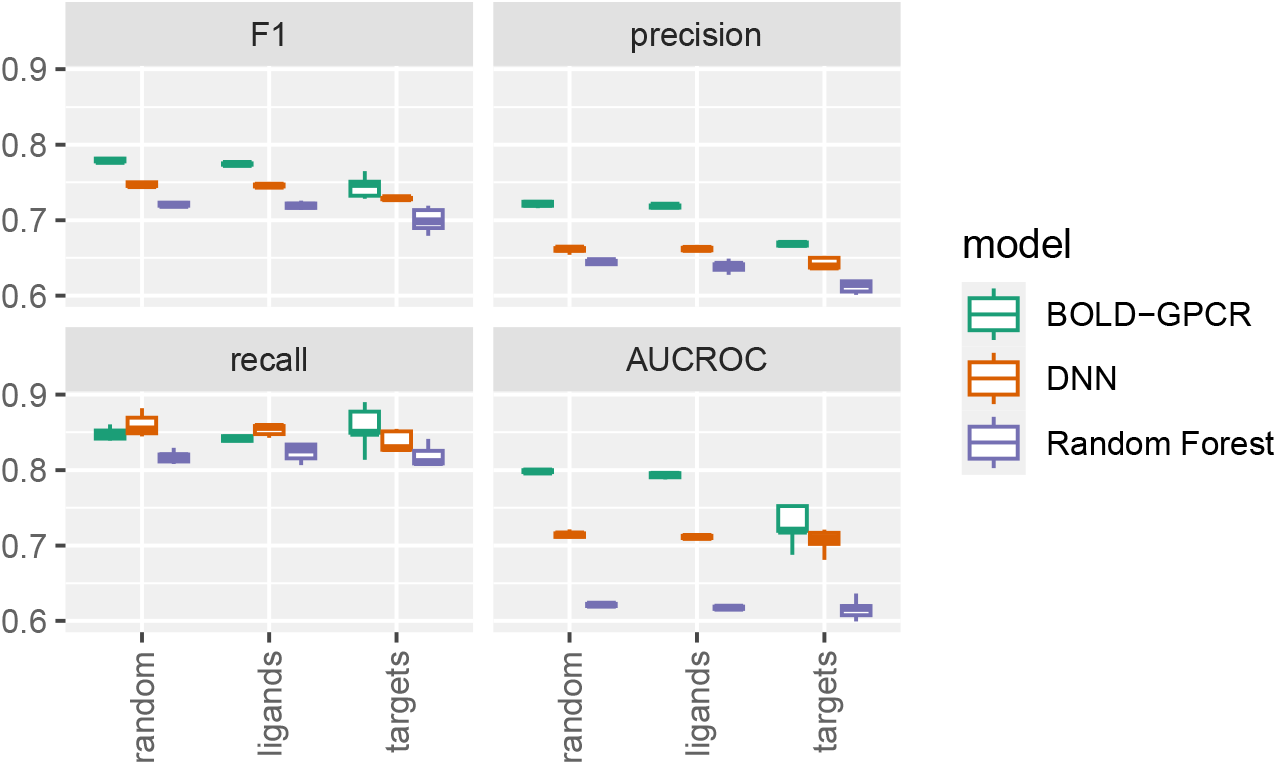
Model performance on the complete Class A GPCR dataset. Metrics are calculated on the 20% validation set. The box plots show results from the BOLD-GPCRs’ architecture (green), a conventional DNN (orange), and a Random Forest model (purple).

Compared to simpler models that do not leverage GPCR embeddings via a LLM, specifically a conventional DNN and a Random Forest classifier trained on ligand molecular properties and one-hot encoded receptor labels, the BOLD-GPCRs’ architecture significantly boosts precision on random splits from an average of 0.66 to 0.75 or 0.64 to 0.72, respectively. Similar improvements are observed in F1 score and AUCROC (Figure 2 and Table S3). Notably, BOLD-GPCRs’ model maintains a high average recall of 0.85, closely matching the baseline DNN classifier at 0.86 and slightly outperforming the Random Forest classifier at 0.82.

This performance trend holds in a more stringent evaluation setting where ligands in the validation set are excluded from the training set (Table S3). In this setting, the BOLD-GPCRs’ architecture achieves an average precision of 0.72, outperforming the DNN (0.66) and Random Forest (0.64) classifiers, while preserving a strong average recall of 0.84, compared to 0.86 for the DNN and 0.82 for the Random Forest. We assessed the model on the ligand predictions on wild-type targets in the training data. Despite the dataset being heavily skewed towards active datapoints (93% positives), the model’s performance was hindered by the limited size of the mutation data. While the model achieved a high precision of 0.88 (comparable to the one on wild type), its recall was low at 0.43, indicating it failed to identify more than half of the actual positive cases. By adding an additional epoch of few-shot fine-tuning, the model’s ability to correctly identify positive cases improved considerably. The recall improved to 0.96, while the precision was largely maintained at 0.85. This demonstrates the model’s ability to generalize effectively with minimal targeted training.

### BOLD-GPCRs’ Architecture Performance is Independent of Ligand Count or Receptor Structure Availability

We next examined the performance of the BOLD-GPCRs architecture within individual class A GPCR subfamilies (Figure 3).

**Figure 3.**
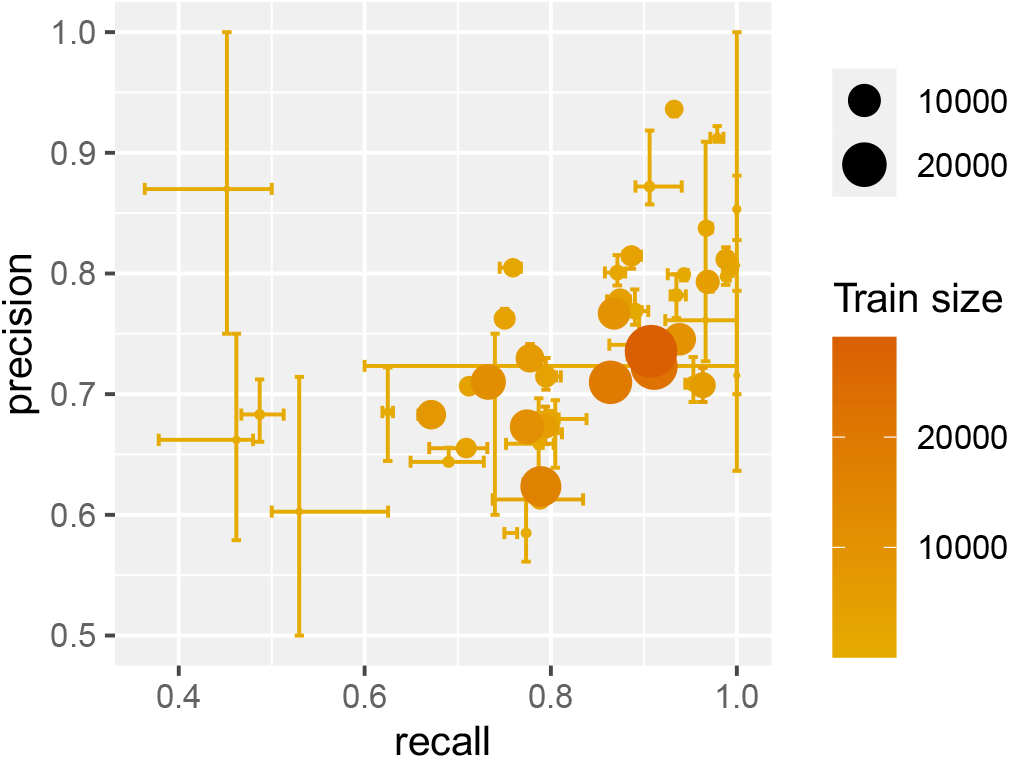
Precision and recall values for each class A GPCR subfamily across random validation split sets. Error bars represent the 25th and 75th quantiles from cross-validation. The color and size of each point correspond to the size of the respective training set.

Table S4 lists GPCRs for which the BOLD-GPCRs’ architecture yields an average precision or recall below 0.63, while Table S5 highlights GPCRs for which the model achieves both average precision and recall above 0.80 on the test set. Notably, strong model performance does not necessarily correlate with the number of available ligands or receptor structures. Although 13 of the 23 receptors with the lowest average precision or recall have fewer than 500 ligands, 10 have more ligands than two of the receptors with the highest model performance (compare Table S4 with Table S5). Figure 4 further illustrates the precision attained by BOLD-GPCRs’ architecture across individual class A GPCRs, mapped onto a phylogenetic tree based on sequence similarity.

**Figure 4.**
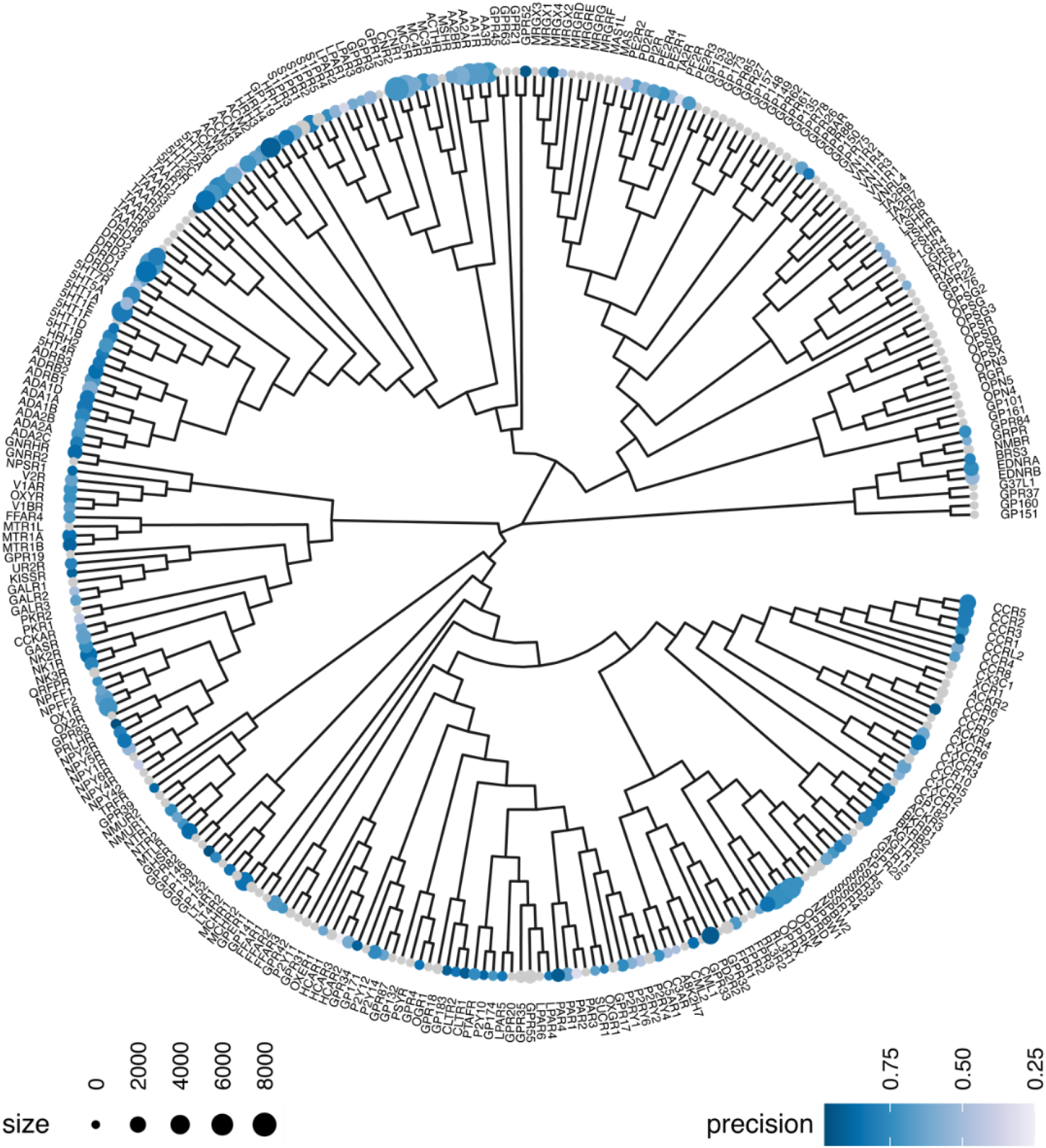
A phylogenetic tree depicting sequence similarity among class A GPCRs, with each node sized according to the number of ligands in the training set. Node color indicates the precision achieved by the BOLD-GPCRs architecture on the test set, ranging from light blue (low precision) to dark blue (high precision).

To evaluate the predictive performance of BOLD-GPCRs’ architecture on GPCRs lacking known ligands, commonly referred to as orphan GPCRs, we explored whether specific dataset characteristics could effectively forecast model accuracy. In particular, we analyzed the relationship between model performance and three target-specific variables: (1) the effective training set size (*ñ*_*j*_), (2) the effective chemical diversity of ligands in the training set 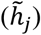, and (3) the degree of class imbalance in the effective training set 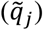. While each variable individually exhibited only a weak correlation with performance metrics (see Table S6), their combined influence was effectively captured using a simple Random Forest regression model. This model yielded a correlation coefficient of 0.93, indicating that although no single feature is strongly predictive on its own, the interplay among these variables provides a reliable foundation for estimating BOLD-GPCRs’ architecture performance on orphan GPCRs.

### BOLD-GPCRs’ Architecture Trained Exclusively on Ligand Features Accurately Predicts GPCR Ligand Bioactivity For Unseen Receptors, With Only a Modest Performance Drop

To assess the generalizability of the BOLD-GPCRs’ architecture to unseen targets, we evaluated its performance on GPCRs excluded from the training set. Using validation splits that withheld all targets (see Fig. 2), the BOLD-GPCRs’ architecture again outperformed both baseline models, achieving higher average precision (0.66 versus 0.64 and 0.62 for the DNN and Random Forest classifiers, respectively) and recall (0.86 versus 0.84 and 0.81 for the DNN and Random Forest classifiers, respectively).

To further test the BOLD-GPCRs’ architecture under more stringent conditions, we designed a challenge scenario in which the entire opioid receptor subfamily, comprising the three primary receptors (OPRD, OPRK, and OPRM) and the nociceptin receptor (OPRX), was excluded from the training set. In this scenario, the architecture was still able to predict the bioactivities of ligands for the excluded targets with acceptable values of precision and recall, despite a significant performance loss. When trained on the full dataset, the model achieved an average precision of 0.78 and recall of 0.89 across these four targets. In contrast, when all opioid receptor ligands were removed from the training set, BOLD-GPCRs’ architecture achieved an average precision of 0.68 and recall of 0.67. The values for the individual primary receptors also reflected this decline, with average precision values of 0.62, 0.69, and 0.69 (down from 0.82, 0.71, and 0.80), and average recall values of 0.78, 0.59, and 0.62, (down from 0.89, 0.89, and 0.86) for OPRD, OPRK, and OPRM, respectively. For OPRX, the average precision and recall values were 0.81 (down from 0.86) and 0.89 (down from 1.00), respectively.

### BOLD-GPCRs Uses Attention Metrics to Highlight Key Residues in Receptor Activation

BOLD-GPCRs’ transformer architecture includes global attention heads that identify residues contributing to changes in global protein embedding, features ultimately used to predict ligand activity at a given receptor. Each attention head *ℓ* generates a query vector *q*_*ℓ*_ from the global embedding *x*, while each amino acid embedding is transformed into a key vector *k*_*iℓ*_. The interaction between *q*_*ℓ*_ and *k*_*iℓ*_, computed as from the inner product ⟨*q*_*ℓ*_, *k*_*iℓ*_⟩, yields a set of normalized attention weights *z*_*iℓ*_· For each residue *i*, we report the average attention score *z*_*i*_ = ∑_*ℓ*_ *z*_*iℓ*_/24, calculated across 4 attention heads and 6 transformer blocks for selected receptor targets. We focused on targets with larger training datasets across major GPCR subfamilies, including peptide receptors (e.g., opioid, angiotensin), aminergic receptors (e.g., 5HT, dopamine, acetylcholine, adrenergic, histamine), protein receptors (e.g., chemokine), nucleotide receptors (e.g., adenosine), and lipid receptors (e.g., cannabinoid). To enable consistent residue-level comparisons, we calculated average attention scores 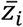 using human GPCR sequences and selected residues according to their GPCRdb generic numbering.^36^ Residues ranking in the top 10% of average attention for at least one GPCR are shown in Figure 5.

**Figure 5.**
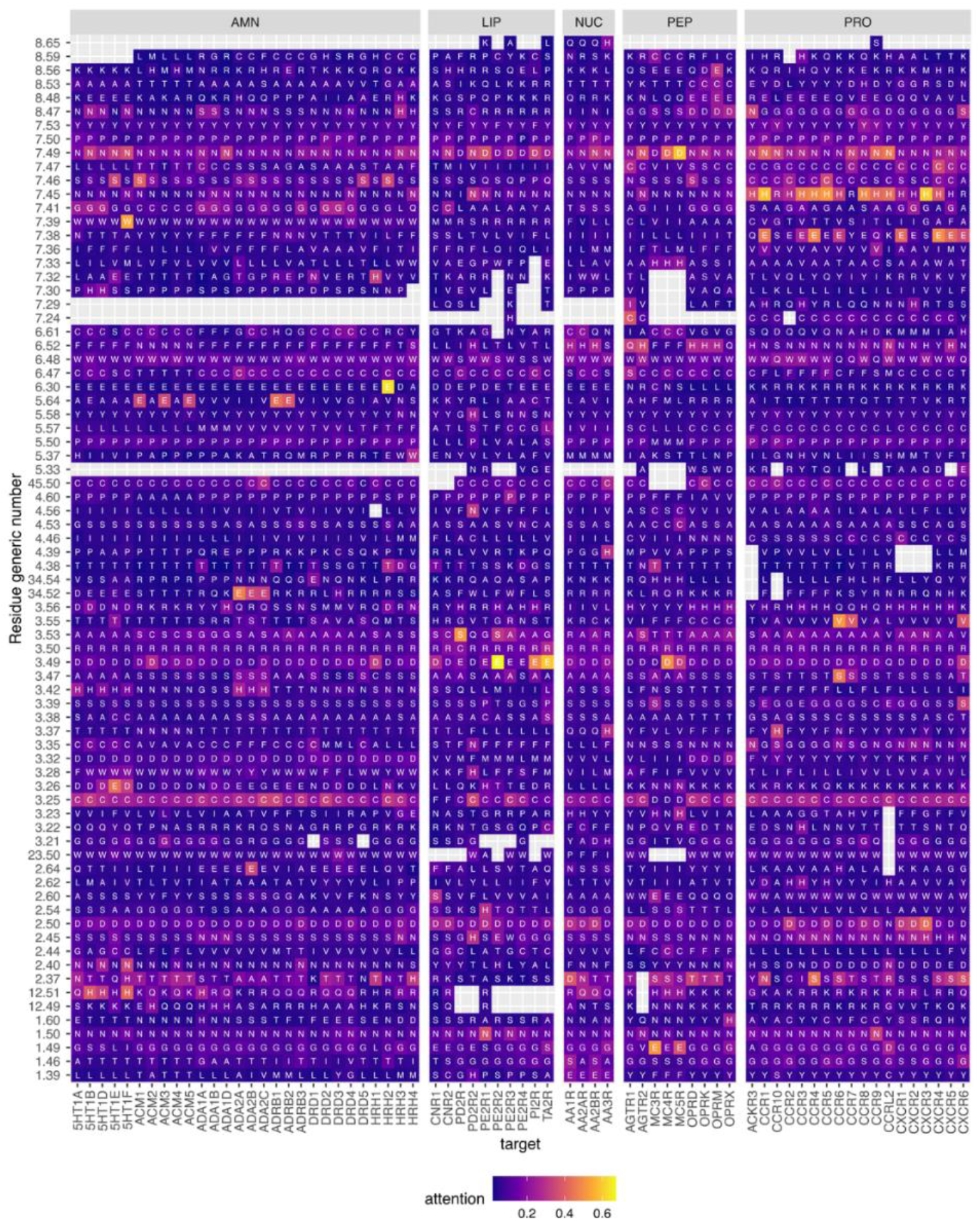
Average attention values for residues with assigned generic numbers and ranked within the top 90^th^ percentile of attention scores. Only receptors from the main GPCR subfamilies with the largest training sets are shown: aminergic receptors (AMN), lipid receptors (LIP), nucleotide receptors (NUC), peptide receptors (PEP), and protein receptors (PRO).

To investigate the relationship between attention scores and residue conservation, we computed the Shannon entropy of amino acid distributions at each alignment position using GPCRdb structural alignments. Entropy at position *i* was calculated as: *H*_*i*_ = − ∑_*s*_ *p*_*is*_log_2_(*p*_*is*_) where *pis* denotes the frequency of amino acid *s* at position *i*. The change in average attention score for positions with different levels of conservation within the top 90^th^ percentile of the attention distribution is displayed in Figure 6 and Table S7. As expected, residues with low entropy, indicative of high conservation, tend to receive higher average attention scores. However, several positions with relatively high entropy (i.e., less conserved) also exhibit strong attention scores, suggesting functional significance beyond sequence conservation. For example, residue 12.51 in ICL1 (H ≈ 2.21 bits, average attention ≈ 12%), its neighbor 2.37 at the cytoplasmic end of TM2 (H ≈ 2.50 bits, attention ≈ 16%), and residue 3.53 at the G protein interface (H ≈ 2 bits, attention ≈ 14%) all display notable attention despite higher entropy levels. Several residues within orthosteric ligand binding pockets also stand out with high attention scores. These residues include: 3.32 (the conserved aspartic acid in aminergic and some peptide receptors), 2.50 (a sodium ion-coordinating site), 2.54, 2.60, 3.35, 3.37, and residues within the CWxP motif of TM6 (6.47, 6.48), as well as 6.52, 7.38, 7.39, 7.41, and 7.46.

**Figure 6.**
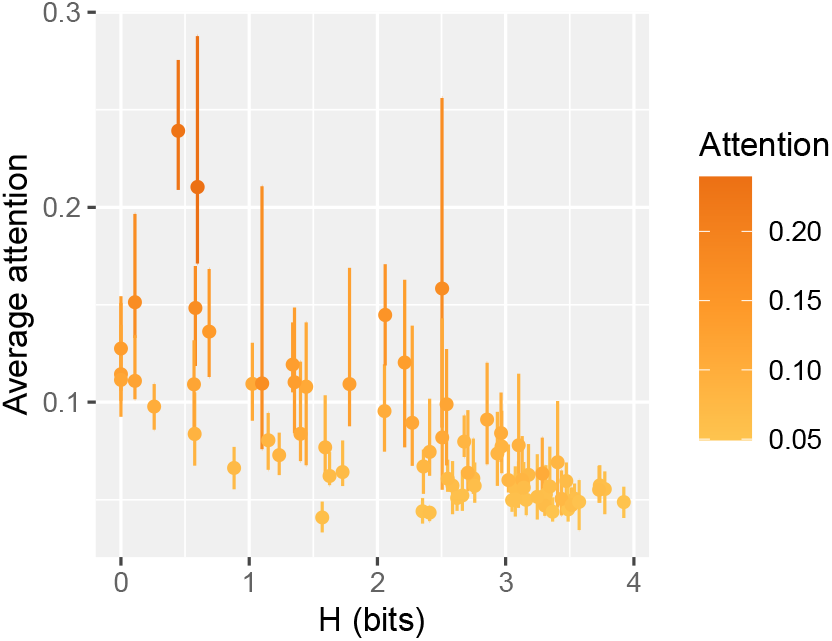
Relationship between residue conservation and average attention scores in Class A GPCRs. The average attention assigned to each residue is plotted against its conservation score derived from the Class A GPCR alignment in GPCRdb. Each data point represents a single residue, with error bars indicating the 25^th^ and 75^th^ percentiles across 24 attention values per residue.

Regions implicated in allosteric modulation are also characterized by substantial attention scores. These include the D(E)RY motif in TM3 (e.g., 3.49, 3.50), the NPxxY motif in TM7 (e.g., 7.49, 7.50, 7.53), and residue 1.46, located in the hydrophobic core and associated with helix packing and activation-related conformational changes.

Residues within ICL2 or in it close proximity, such as 3.55, 3.56, 34.52, 34.54, 4.38, and 4.39, also rank in the top 10% of attention scores, underscoring their known roles in G protein interaction and activation modulation.^37, 38^ Similarly, several residues in helix 8, including 8.47, 8.48, 8.53, 8.56, 8.59, and 8.65, exhibit high attention scores, consistent with their known involvement in arrestin recruitment.^39, 40^

Taken together, this attention analysis suggests that the learned protein embeddings prioritize residues located in key receptor regions critical for ligand recognition, activation, and effector coupling.

### BOLD-GPCRs Features a User-Friendly Web Interface

We developed an intuitive, web-based platform that provides seamless access to the pretrained BOLD-GPCRs’ architecture. As illustrated in Figure 7, the landing page offers a clean, centralized interface for user interaction. Through this interface, users can input class A GPCR sequences in FASTA^41^ format and ligand structures in SMILES^24^ format either by directly entering them into input fields or by uploading CSV files. The platform then returns predictions of ligand bioactivity. To support high-throughput applications, the interface allows the simultaneous submission of multiple class A GPCR sequences and up to 1,000 ligands. Users can opt to compute ligand bioactivity predictions for all possible receptor sequence-ligand combinations, enabling efficient assessment of selectivity and potential off-target effects, or restrict predictions to specified sequence–ligand pairs. Designed for flexibility and ease of use, the platform offers the following options: (a) Select from a curated list of human class A GPCR sequences; (b) Upload custom GPCR sequences and/or ligand libraries. A sample GPCR-ligand pair is also available for demonstration purposes.

**Figure 7.**
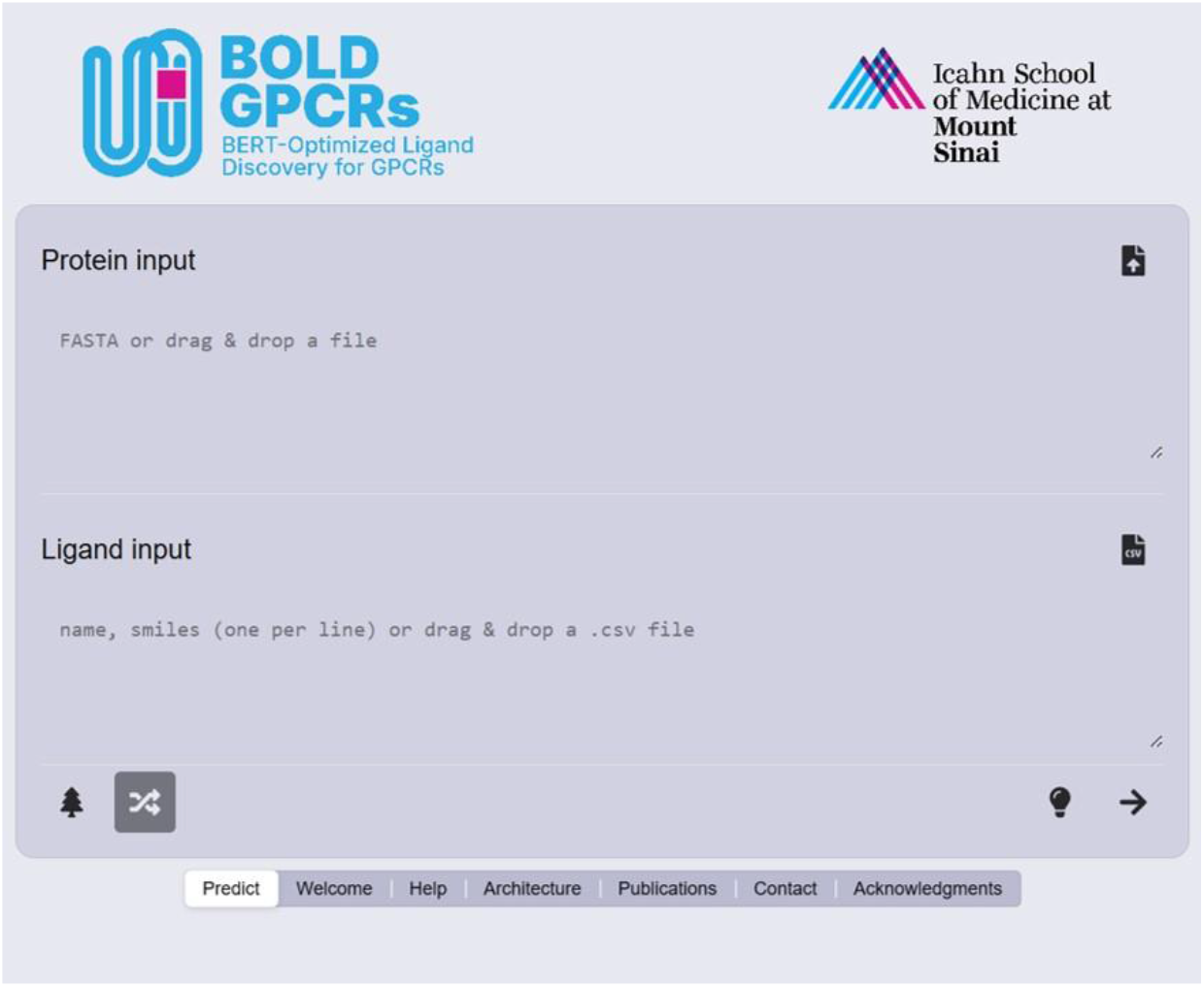
BOLD-GPCRs’ landing page. The interface features a central input module where users can specify class A GPCRs and ligands, either by selecting from preset lists or by uploading custom files.

Prediction results are displayed directly on the website in a downloadable table listing GPCR names, sequences, ligand names, and SMILES strings. Internal benchmarks indicate that the server can process approximately 1,000 bioactivity predictions per minute. The backend pipeline utilizes RDKit^25^ for ligand feature generation and TensorFlow^42^ for bioactivity prediction. The server is hosted locally and its use is intended solely for research and informational purposes.

## DISCUSSION

The enhanced performance of BOLD-GPCRs relative to traditional DNN and Random Forest models underscores the value of protein embeddings derived from large language models in capturing the intricate functional landscape of GPCR-ligand interactions. Unlike methods that depend solely on receptor identity or molecular fingerprints, BOLD-GPCRs demonstrates a robust ability to generalize across diverse GPCR subfamilies, even when specific ligands or entire receptor subtypes are excluded from the training set. While precision improves consistently across datasets, recall remains comparable to that of baseline models. This pattern may indicate that the BOLD-GPCRs’ architecture is particularly adept at minimizing false positives rather than broadening the recovery of active compounds. Such a tradeoff could stem from the inductive biases introduced by transformer-based attention layers or from the composition of actives in the training data. Future work might explore whether more advanced ligand embedding techniques, such as those based on molecular graphs or chemical language models, could improve the precision-recall balance. Additionally, alternative communication strategies between ligand and protein representations, including cross-attention mechanisms, could be explored. We also plan to evaluate ensemble modeling and multi-task learning approaches as potential avenues to enhance recall without compromising precision.

The interpretability afforded by attention scores offers valuable insights into the inner workings of the BOLD-GPCRs’ model, prompting intriguing questions about the nature of the encoded embeddings. High attention scores frequently localize to conserved functional motifs, such as the DRY and NpxxY motifs, sodium-binding residues, and key positions within ICL2 and helix 8, indicating that the model is indeed learning key mechanistic features underlying GPCR signaling. These patterns may emerge during pre-training, fine-tuning, or both.

Because the BOLD-GPCRs’ architecture integrates residue-level embeddings, it is also sensitive to localized sequence perturbations such as mutations, providing an initial step toward predicting how specific variants may modulate receptor behavior or ligand response. Intriguingly, high-attention residues with relatively high entropy (e.g., 12.51, 2.37, 3.53) indicate that attention is not simply a proxy for conservation. Instead, it appears that the model discerns context-specific functional relevance that escapes traditional sequence-based metrics. This observation aligns with recent findings that language models can implicitly learn structural and functional features solely from sequence data, thus reinforcing their potential utility in biological inference.

Despite its good performance, the BOLD-GPCRs framework has several limitations worth noting. Experimental validation of residues with high-attention scores, particularly those not previously identified as functionally critical, will be necessary to determine whether the model identifies novel biological insights or is instead overfitting to patterns present in the data. Additionally, although the web server facilitated practical application, its predictive reliability differs among receptor classes with limited ligand data or atypical pharmacological profiles. Expanding the training datasets with additional experimental or high-confidence synthetic data, and incorporating three-dimensional structural information, could further improve model performance and generalizability.

Finally, evaluating the model’s predictive ability on receptors similar to those excluded from the training set, such as olfactory Class A targets and non-Class A GPCRs, as well as exploring more nuanced pharmacological tasks, such as separating potency from efficacy or introducing multiple signaling endpoints, may provide further insights into the scope and limitations of this approach.

## CONCLUSIONS

Our findings indicate that combining ligand descriptors with protein-language model embeddings yields both high recall and robust precision in predicting GPCR ligand bioactivities. For most targets, the model achieves precision above 0.60 and recall above 0.75. Importantly, its performance remains strong even on receptors excluded from training, demonstrating the model’s capacity to generalize beyond memorized ligand–receptor pairs and infer functional relationships from sequence and ligand information alone. The built-in interpretability of the attention mechanisms further underscores the model’s value: it consistently highlights residues implicated in receptor activation, G protein coupling, and arrestin recruitment, despite having access only to receptor sequence and ligand input. This makes BOLD-GPCRs especially well-suited for exploring understudied receptors with limited experimental data.

By bridging protein-language models with ligand-based embeddings, BOLD-GPCRs represents a step forward in data-driven ligand discovery for GPCRs. Future extensions could incorporate receptor-specific structural or context-dependent features, broaden applicability to other receptor families, or support studies in polypharmacology and pathway-selective signaling. Overall, BOLD-GPCRs provides a scalable, interpretable, and effective framework for virtual screening across the GPCR landscape.

## DATA AND MATERIAL AVAILABILITY

The web tool is accessible at https://bold-gpcrs.filizolalab.org/. Processed training datasets are available on Zenodo (DOI: 10.5281/zenodo.16740418).

## ACKNOWLEDGMENTS

This work was funded by National Institutes of Health grant DA059420. Computations were supported in part through the computational resources and staff expertise provided by Scientific Computing at the Icahn School of Medicine at Mount Sinai, the Clinical and Translational Science Awards (CTSA) grant UL1TR004419 from the National Center for Advancing Translational Sciences, and the Office of Research Infrastructure of the National Institutes of Health under award number S10OD026880 and S10OD030463. The authors thank the Scientific Computing team, with special appreciation to Wei Guo, for their assistance with application security assessments. The content is solely the responsibility of the authors and does not necessarily represent the official views of the National Institutes of Health.

