## Supplemental Tables for "BOLD-GPCRs: A Transformer-Powered App for Predicting Ligand Bioactivity and Mutational Effects Across Class A GPCRs"

Table S1 ........................................................................................................pg. 2

Table S2 ........................................................................................................pg. 3

Table S3 ........................................................................................................pg. 4

Table S4 ........................................................................................................pg. 5

Table S5 ........................................................................................................pg. 6

Table S6. ......................................................................................................pg. 7

Table S7. ......................................................................................................pg.8

**Table S1.** Summary of the complete training set of class A GPCRs, displaying the total number of data points used.

| | Total bioactivities | Unique ligands | Unique receptors |
|---|---|---|---|
| Active | 214,816 | 138,232 | 566 |
| Inactive | 168,822 | 74,953 | 531 |
| Total | 383,638 | 180,340 | 615 |

**Table S2.** The total number of data points, unique ligands, and unique receptors for each class A GPCR subfamily, presented in slash-separated format.

| Family Name | Inactive | Active | Total |
|---|---:|---:|---:|
| 5-Hydroxytryptamine receptors | 17,194/8,574/42 | 28,983/18,528/46 | 46,177/23,915/49 |
| Acetylcholine receptors (muscarinic) | 11,058/5,137/18 | 7,620/4,196/17 | 18,678/8,499/18 |
| Adenosine receptors | 16,119/8,614/21 | 16,276/9,342/23 | 32,395/14,272/23 |
| Adrenoceptors | 13,855/5,107/37 | 11,750/5,965/37 | 25,605/9,176/42 |
| Angiotensin receptors | 2,588/2,471/9 | 3,446/2,638/10 | 6,034/4,788/10 |
| Apelin receptor | 254/251/2 | 589/498/2 | 843/746/2 |
| Bile acid receptor | 480/400/3 | 700/522/3 | 1,180/846/3 |
| Bombesin receptors | 363/288/10 | 1,037/676/11 | 1,400/865/11 |
| Bradykinin receptors | 1,795/1,479/7 | 1,533/1,271/10 | 3,328/2,640/10 |
| Cannabinoid receptors | 9,521/7,591/6 | 11,262/8,392/6 | 20,783/12,270/6 |
| Chemokine receptors | 4,425/3,917/36 | 9,807/9,139/46 | 14,232/12,632/49 |
| Cholecystokinin receptors | 4,308/2,952/8 | 3,152/2,246/9 | 7,460/4,197/9 |
| Class A Orphans | 3,080/2,689/42 | 4,738/4,304/38 | 7,818/6,865/45 |
| Dopamine receptors | 14,489/8,519/21 | 18,816/11,894/20 | 33,305/17,430/22 |
| Endothelin receptors | 3,021/2,288/8 | 2,987/2,078/9 | 6,008/3,524/9 |
| Free fatty acid receptors | 1,428/1,185/11 | 1,816/1,486/11 | 3,244/2,413/13 |
| Ghrelin receptor | 1,656/1,656/3 | 1,722/1,692/6 | 3,378/3,344/6 |
| Gonadotrophin-releasing hormone receptors | 506/476/2 | 2,055/1,763/5 | 2,561/2,150/5 |
| Histamine receptors | 7,285/4,283/15 | 9,401/7,392/15 | 16,686/10,704/16 |
| Hydroxycarboxylic acid receptors | 570/498/6 | 592/538/5 | 1,162/1,000/6 |
| Leukotriene receptors | 1,160/1,026/11 | 1,791/1,626/11 | 2,951/2,554/12 |
| Lysophospholipid (LPA) receptors | 315/216/8 | 518/345/10 | 833/483/10 |
| Lysophospholipid (S1P) receptors | 3,113/2,199/5 | 3,724/2,774/10 | 6,837/3,770/10 |
| Melanin-concentrating hormone receptors | 964/920/4 | 3,355/3,096/6 | 4,319/3,912/6 |
| Melanocortin receptors | 6,112/3,408/14 | 5,196/3,155/10 | 11,308/5,742/14 |
| Melatonin receptors | 425/306/3 | 1,819/1,056/5 | 2,244/1,246/5 |
| Motilin receptor | 1,276/1,276/2 | 324/316/2 | 1,600/1,590/2 |
| Neuropeptide Y receptors | 3,077/2,141/9 | 2,584/2,290/8 | 5,661/4,131/9 |
| Neurotensin receptors | 790/716/5 | 445/334/5 | 1,235/1,009/5 |
| Opioid receptors | 15,159/8,455/16 | 22,706/10,922/18 | 37,865/15,609/18 |
| Orexin receptors | 2,707/2,084/4 | 5,355/3,332/6 | 8,062/4,168/6 |
| P2Y receptors | 1,763/1,375/17 | 2,048/1,845/16 | 3,811/3,019/18 |
| Platelet-activating factor receptor | 489/488/3 | 972/961/4 | 1,461/1,446/4 |
| Prostanoid receptors | 6,062/4,152/24 | 7,031/6,017/27 | 13,093/9,185/27 |
| Proteinase-activated receptors | 674/665/4 | 1,436/1,436/3 | 2,110/2,078/4 |
| Somatostatin receptors | 2,207/976/11 | 2,930/1,559/11 | 5,137/1,930/13 |
| Tachykinin receptors | 1,607/1,205/13 | 4,713/3,886/15 | 6,320/4,788/15 |
| Trace amine receptor | 733/672/7 | 2,418/1,521/5 | 3,151/2,106/7 |
| Urotensin receptor | 216/209/4 | 673/554/4 | 889/733/5 |
| Vasopressin and oxytocin receptors | 4,072/2,483/10 | 3,703/2,424/12 | 7,775/4,099/12 |
| Other | 1,906/1,769/50 | 2,793/2,280/49 | 4,699/3,927/59 |

**Table S3.** Comparison of BOLD-GPCRs, DNN, and random forest models in terms of average F1 score, precision, recall, and AUCROC, with the 25$^{th}$ and 75$^{th}$ quantiles provided in parentheses.

| **Split** | **Model** | **F1** | **precision** | **recall** | **AUCROC** |
|---|---|---|---|---|---|
| random | BOLD-GPCR | 0.78 (0.78,0.78) | 0.72 (0.72,0.72) | 0.85 (0.84,0.85) | 0.80 (0.80,0.80) |
| random | DNN | 0.75 (0.75,0.75) | 0.66 (0.66,0.66) | 0.86 (0.85,0.87) | 0.71 (0.71,0.72) |
| random | Random Forest | 0.72 (0.72,0.72) | 0.64 (0.64,0.65) | 0.82 (0.81,0.82) | 0.62 (0.62,0.62) |
| ligands | BOLD-GPCR | 0.77 (0.77,0.78) | 0.72 (0.72,0.72) | 0.84 (0.84,0.84) | 0.79 (0.79,0.80) |
| ligands | DNN | 0.75 (0.74,0.75) | 0.66 (0.66,0.66) | 0.86 (0.85,0.86) | 0.71 (0.71,0.71) |
| ligands | Random Forest | 0.72 (0.72,0.72) | 0.64 (0.63,0.64) | 0.82 (0.82,0.83) | 0.62 (0.62,0.62) |
| targets | BOLD-GPCR | 0.74 (0.73,0.75) | 0.66 (0.66,0.67) | 0.86 (0.85,0.88) | 0.73 (0.72,0.75) |
| targets | DNN | 0.73 (0.73,0.73) | 0.64 (0.64,0.65) | 0.84 (0.83,0.85) | 0.71 (0.70,0.72) |
| targets | Random Forest | 0.70 (0.69,0.71) | 0.62 (0.61,0.62) | 0.81 (0.81,0.83) | 0.62 (0.61,0.62) |

**Table S4.** A list of GPCRs for which the BOLD-GPCRs architecture achieves suboptimal performance on the test set, as indicated by average precision or recall values below 0.63. For each receptor, the table reports the total number of known ligands and the number of available high-resolution experimental structures.

| Label | Receptor | Structures | Ligands | F1 | Recall | Precision |
|---|---|---|---|---|---|---|
| ACM5 | Muscarinic acetylcholine receptor M5 | 1 | 1097 | 0.53 | 0.84 | 0.38 |
| NPY1R | Neuropeptide Y receptor type 1 | 6 | 976 | 0.43 | 0.36 | 0.55 |
| MC3R | Receptor for Melanocyte-stimulating hormone (MSH) andAdrenocorticotropic Hormone (ACTH) | 1 | 749 | 0.33 | 0.23 | 0.57 |
| HRH2 | Histamine H2 receptor | 5 | 684 | 0.63 | 0.51 | 0.82 |
| NPY2R | Neuropeptide Y receptor type 2 | 5 | 666 | 0.20 | 0.11 | 0.80 |
| 5HT5A | 5-hydroxytryptamine receptor 5A | 5 | 664 | 0.36 | 0.28 | 0.51 |
| S1PR4 | Sphingosine 1-phosphate receptor 4 | 1 | 663 | 0.60 | 0.66 | 0.56 |
| PE2R2 | Prostaglandin E2 receptor EP2 subtype | 3 | 655 | 0.37 | 0.33 | 0.42 |
| MSHR | Melanocyte-stimulating hormone | 4 | 572 | 0.64 | 0.65 | 0.63 |
| DRD5 | $D_5$ or D(1B) dopamine receptor | 1 | 562 | 0.59 | 0.79 | 0.47 |
| NPBW1 | Neuropeptides B/W receptor type 1 | 0 | 377 | 0.55 | 0.44 | 0.73 |
| P2RY2 | Nucleotide receptors P2Y | 0 | 343 | 0.27 | 0.19 | 0.46 |
| P2RY6 | P2Y purinoceptor 6 | 0 | 320 | 0.25 | 0.16 | 0.57 |
| GALR3 | Galanin receptor type 3 | 0 | 256 | 0.44 | 0.33 | 0.67 |
| PF2R | Prostaglandin F2-alpha receptor. | 8 | 241 | 0.27 | 0.23 | 0.32 |
| FFAR2 | Free fatty acid receptor 2 | 5 | 221 | 0.40 | 0.28 | 0.71 |
| C3AR | C3a anaphylatoxin chemotactic rec. | 9 | 164 | - | - | 0.48 |
| PKR1 | Prokineticin receptors | 0 | 162 | - | - | 0.44 |
| PAR2 | Proteinase-activated receptors PAR2 | 3 | 131 | 0.47 | 0.89 | 0.32 |
| LPAR2 | Lysophosphatidic acid receptor 2 | 1 | 128 | 0.43 | 0.55 | 0.35 |
| HCAR3 | Hydroxycarboxylic acid receptor 3 | 4 | 116 | 0.29 | 0.33 | 0.25 |
| GALR2 | Galanin receptors GAL2 receptor | 4 | 95 | 0.29 | 0.20 | 0.50 |
| NPY4R | Neuropeptide Y receptors Y4 receptor | 1 | 82 | 0.18 | 0.12 | 0.40 |

**Table S5.** A list of GPCRs for which the BOLD-GPCRs architecture achieves both an average precision and average recall exceeding 0.80 on the test set. For each receptor, the table reports the total number of known ligands and the number of available high-resolution experimental structures.

| Label | Receptor | Structures | Ligands | F1 | recall | precision |
|---|---|---|---|---|---|---|
| BKRB2 | B2 bradykinin receptor | 3 | 499 | 0.92 | 0.99 | 0.85 |
| TAAR1 | Trace amine-associated rec. 1 | 2 | 578 | 0.93 | 0.92 | 0.94 |
| DRD3 | D3 receptor | 6 | 5282 | 0.92 | 1.00 | 0.85 |
| MTR1A | Melatonin receptor type 1A | 8 | 1105 | 0.92 | 0.99 | 0.86 |

**Table S6.** Pearson correlations between model performance metrics (F1 score, precision, and recall) and various properties of the effective training set, including the number of active, inactive, and total ligands; the chemical heterogeneity of active, inactive, and total ligands; and the degree of class imbalance.

| | Inactives | Actives | Total | Het. Act. | Het. Inact. | Het. Tot. | Imbalance |
|---|---|---|---|---|---|---|---|
| F1 | 0.057 | 0.068 | 0.063 | 0.013 | -0.122 | 0.016 | 0.061 |
| precision | -0.026 | -0.015 | -0.02 | 0.174 | -0.098 | 0.075 | 0.247 |
| recall | 0.032 | 0.043 | 0.039 | -0.021 | -0.022 | 0.071 | 0.045 |

**Table S7.** Average attention value and Shannon entropy for positions with attention values in the top 10%.

| Generic Number | Attention | H |
|---|---|---|
| 7.49 | 0.24 (0.17,0.29) | 0.60 |
| 3.25 | 0.23 (0.21,0.28) | 0.45 |
| 2.37 | 0.17 (0.09,0.26) | 2.50 |
| 2.50 | 0.17 (0.13,0.2) | 0.11 |
| 3.49 | 0.17 (0.12,0.17) | 0.58 |
| 7.45 | 0.17 (0.08,0.21) | 1.10 |
| 3.53 | 0.16 (0.12,0.17) | 2.06 |
| 1.50 | 0.14 (0.11,0.15) | - |
| 6.48 | 0.14 (0.11,0.17) | 0.69 |
| 1.49 | 0.13 (0.11,0.14) | 1.34 |
| 12.51 | 0.13 (0.08,0.16) | 2.21 |
| 2.45 | 0.13 (0.09,0.17) | 1.78 |
| 45.50 | 0.13 (0.1,0.15) | - |
| 7.46 | 0.13 (0.08,0.15) | 1.35 |
| 3.50 | 0.12 (0.1,0.13) | 0.11 |
| 6.52 | 0.12 (0.07,0.13) | 2.54 |
| 7.24 | 0.12 (0.09,0.13) | 0.57 |
| 7.50 | 0.12 (0.09,0.14) | - |
| 3.47 | 0.11 (0.07,0.12) | 1.40 |
| 5.50 | 0.11 (0.09,0.13) | 1.02 |
| 6.47 | 0.11 (0.07,0.14) | 1.44 |
| 7.41 | 0.11 (0.07,0.14) | 2.27 |
| 8.47 | 0.11 (0.06,0.14) | 2.51 |
| 1.46 | 0.1 (0.07,0.12) | 2.06 |
| 3.35 | 0.1 (0.07,0.11) | 2.96 |
| 7.38 | 0.1 (0.05,0.08) | 3.29 |
| 7.47 | 0.1 (0.07,0.12) | 2.85 |
| 7.53 | 0.1 (0.09,0.11) | 0.26 |
| 23.50 | 0.09 (0.07,0.11) | 0.57 |
| 3.26 | 0.09 (0.05,0.1) | 2.70 |
| 3.32 | 0.09 (0.06,0.1) | 2.41 |
| 3.39 | 0.09 (0.06,0.08) | 1.23 |
| 3.56 | 0.09 (0.06,0.11) | 3.10 |
| 2.40 | 0.08 (0.05,0.08) | 2.36 |
| 3.21 | 0.08 (0.06,0.1) | 1.59 |
| 3.28 | 0.08 (0.06,0.1) | 2.97 |
| 3.42 | 0.08 (0.05,0.08) | 3.02 |
| 3.55 | 0.08 (0.05,0.08) | 3.13 |

| Generic Number | Attention | H |
|---|---|---|
| 4.60 | 0.08 (0.07,0.09) | 1.15 |
| 5.57 | 0.08 (0.07,0.09) | 2.68 |
| 5.64 | 0.08 (0.04,0.06) | 3.43 |
| 6.61 | 0.08 (0.05,0.1) | 3.40 |
| 7.39 | 0.08 (0.06,0.1) | 2.93 |
| 2.54 | 0.07 (0.05,0.08) | 2.75 |
| 2.60 | 0.07 (0.05,0.08) | 3.18 |
| 2.64 | 0.07 (0.05,0.07) | 3.47 |
| 3.37 | 0.07 (0.05,0.07) | 2.55 |
| 3.38 | 0.07 (0.06,0.08) | 1.73 |
| 34.52 | 0.07 (0.04,0.07) | 3.31 |
| 4.53 | 0.07 (0.06,0.07) | 1.63 |
| 5.58 | 0.07 (0.06,0.08) | 0.88 |
| 7.32 | 0.07 (0.05,0.07) | 3.73 |
| 8.53 | 0.07 (0.04,0.08) | 3.34 |
| 1.39 | 0.06 (0.05,0.07) | 3.10 |
| 1.60 | 0.06 (0.05,0.07) | 3.31 |
| 12.49 | 0.06 (0.05,0.07) | 3.07 |
| 2.44 | 0.06 (0.04,0.07) | 2.58 |
| 2.62 | 0.06 (0.04,0.06) | 3.05 |
| 3.22 | 0.06 (0.05,0.07) | 3.73 |
| 3.23 | 0.06 (0.04,0.06) | 3.54 |
| 4.38 | 0.06 (0.03,0.05) | 1.57 |
| 4.46 | 0.06 (0.04,0.07) | 2.66 |
| 4.56 | 0.06 (0.04,0.06) | 2.62 |
| 5.33 | 0.06 (0.04,0.06) | 3.77 |
| 7.29 | 0.06 (0.04,0.06) | 3.51 |
| 7.30 | 0.06 (0.04,0.06) | 3.16 |
| 7.33 | 0.06 (0.05,0.07) | 3.14 |
| 7.36 | 0.06 (0.05,0.07) | 2.76 |
| 8.48 | 0.06 (0.04,0.07) | 3.25 |
| 8.56 | 0.06 (0.04,0.06) | 3.07 |
| 8.65 | 0.06 (0.04,0.05) | 2.41 |
| 34.54 | 0.05 (0.04,0.06) | 3.49 |
| 4.39 | 0.05 (0.04,0.05) | 3.37 |
| 5.37 | 0.05 (0.04,0.06) | 3.92 |
| 6.30 | 0.05 (0.04,0.05) | 2.35 |
| 8.59 | 0.05 (0.03,0.06) | 3.57 |
